# Extensive variation in germline *de novo* mutations in *Poecilia reticulata*

**DOI:** 10.1101/2023.03.22.533860

**Authors:** Yuying Lin, Iulia Darolti, Wouter van der Bijl, Jake Morris, Judith E. Mank

## Abstract

The rate of germline mutation is fundamental to evolutionary processes, as it generates the variation upon which selection acts. The guppy, *Poecilia reticulata*, is a model of rapid adaptation, however the relative contribution of standing genetic variation versus *de novo* mutation to evolution in this species remains unclear. Here, we use pedigree-based approaches to quantify and characterize *de novo* mutations (DNMs) in three large guppy families. Our results suggest germline mutation rate in the guppy varies substantially across individuals and families. Most DNMs are shared across multiple siblings, suggesting they arose during early zygotic development. DNMs are randomly distributed throughout the genome, and male-biased mutation rate is low, as would be expected from the short guppy generation time. Overall, our study demonstrates remarkable variation in germline mutation rate and provides insights into rapid evolution of guppies.

---

The timing, rate and spectra of germline mutations are fundamental aspects of molecular evolution and the basis of adaptation (Lynch et al., 2016). The rate of mutations is a product of several processes. *De novo* mutations (DNMs) arise from replication errors that damage repair mechanisms fail to correct (Gao et al., 2019; Manuel et al., 2022; Seplyarskiy et al., 2021), and so the fidelity of replication and the efficacy of repair mechanisms are major contributors to the DNM rate. All cell divisions, somatic and germline, experience DNMs, however the rate is far higher in somatic versus germline tissue (Milholland et al., 2017), likely representing different investment in costly repair mechanisms.

Recent sequencing-based approaches to study germline mutation have used pedigrees to characterize germline DNM, and revealed the influence of many factors, including generation time (Bergeron et al., 2023; Carlson et al., 2018; Francioli et al., 2015; Kong et al., 2012; Wang et al., 2022), life history (Bergeron et al., 2023), and number of cell divisions (Ellegren, 2007). However, these factors do not fully explain DNM variation across species (Campbell et al., 2021; Wu et al., 2020; Manuel et al., 2022), and recent work in humans suggests stochastic biological processes and, to a lesser extent, family-specific effects (Goldmann et al., 2021) may be important. Together, these findings indicate that a complex interplay of multiple factors contributing to the variation in the number of DNMs in families.

The guppy, *Poecilia reticulata*, is a model system for rapid ecological adaptation (Gordon et al., 2015; Reznick et al., 1990; Reznick et al., 1997; Whiting et al., 2022) and the swift and convergent patterns of phenotypic adaptation in guppies can arise via two possible genetic mechanisms. It is possible that DNMs underlie the rapid rates of phenotypic adaptation observed in guppies. However, functional mutations are more often deleterious than adaptive (Yoder & Tiley, 2021), and a high germline rate of DNM fostering rapid adaptation could ultimately prove more of a mutational burden than an adaptive boon. Additionally, although somatic mutation rates are strongly anticorrelated with lifespan (Cagan et al., 2022), which is relatively short in guppies, fish are poikilothermic, and we might expect the lower metabolic rate to result in lower rates of DNA damage and therefore mutation rates (Adelman et al., 1988). Consistent with this, DNM rates in fish are slightly lower than homeothermic vertebrates (Bergeron et al., 2023; Feng et al., 2017).

Alternatively, rapid adaptation could be a product of selection on standing genetic variation (Barrett & Schluter, 2008), and consistent with this, natural populations of guppies exhibit extensive genetic and phenotypic polymorphism (Almeida et al., 2020; Lin et al., 2022; Whiting et al., 2021, 2022). The role of DNMs versus standing genetic variation in the rapid adaptation of guppies has important implications to the locus and nature of evolution in this key ecological model, as well as molecular signature we might expect to detect from. Selection on adaptive DNMs will produce a signature of hard sweeps and would most often be associated with a few loci of large effect (Matuszewski et al., 2015; Pritchard et al., 2010). In contrast, selection acting on standing variation is associated with soft sweeps, and will more often results in fixing many alleles of small effect (Hermisson and Pennings, 2017).

Here, we sequenced three large *P. reticulata* families and applied stringent parent-offspring trio-based variant filtering criteria to accurately identify and quantify the number of DNMs, providing insights into the roles of germline mutations in the rapid evolution of this species. We observe a large variation in DNM rate across families. Moreover, our large families, each with ten offspring, allow us to estimate the timing of mutations across germline development, and we observe a large number of DNMs shared across multiple siblings, revealing the mosaic nature of the germline genome through developmental mutations (Goldmann et al., 2019).

## Results

### Identifying DNMs from three guppy pedigrees

We re-sequenced 36 individuals from three guppy pedigrees, each including unrelated parents, five sons and five daughters (Figs. S1-S3), with individual mean sequencing depth ≈ 25X (Table 1 & S1). To limit the impact of genetic differences between the Ensembl female reference genome and the guppy populations in this study, we constructed a high-quality pseudo-genome from a 10X assembly of a female individual taken from the Quare River in Trinidad (see Almeida et al., (2020) for details), the original population from which our lab stock population derives (details see Methods and Material).

**Table 1.** Summary statistics of DNMs in three guppy pedigrees.

| Pedigree | Coverage* | Mapping Rate* | Callable Genome Size* | No. of DNMs (range) | $\mu$, $10^{-8}$ per site per generation (range) |
| --- | --- | --- | --- | --- | --- |
| Fam1 | 26.42X | 99.36% | 432,137,349bp | 5.20<br>(2.00- 9.00) | 0.60<br>(0.23 - 1.27) |
| Fam2 | 24.57X | 99.37% | 428,029,972bp | 24.90<br>(21.00 – 31.00) | 2.90<br>(2.45 - 3.62) |
| Fam3 | 26.10X | 99.31% | 429,364,825bp | 4.80<br>(2.00 – 6.00) | 0.56<br>(0.24 - 0.71) |

\*All data are average across offspring in each pedigree; Details can be found in Tables S1, S6 - S9.

Germline mutations are rare events, and as such are notoriously difficult to differentiate from the various errors introduced from sequencing, read alignments, genotyping and variant filtering. Thus, we applied step-by-step stringent filtering to minimize false discovery rates (Fig. S4 & Table S2). First, we discarded all multiply mapped reads, retaining only uniquely mapped reads for further analysis. Second, we genotyped each parent-offspring trio separately, instead of all offspring in each family together, using two independent genotype callers, GATK v.4.2.6.1 (Van der Auwera & O’Connor, 2020) and BCFtools v.1.16 (Li et al., 2009), and used the intersection of inferred DNMs from both methods. We called sites with Mendelian violations at which parents are homozygous for the same allele while the child is heterozygous as putative DNMs. By applying the same individual-level and site-level filtering criteria, and intersecting the results from the two genotype callers, we obtained an average number of 5.20, 24.90 and 4.80 DNMs across each trio in our three pedigrees (Fam1, Fam2, Fam3) (Table 1 & S1). To calculate the germline mutation rate in each trio, we determined the denominator, the callable genome size, of each individual (see Materials and Methods). After removing repetitive regions and restricting the coverage between > ½ X and < 2X of the individual average sequencing depth, we retained an approximate callable genome size of ≈ 430Mb out of the 720Mb reference genome across individuals (Table 1, S1 & S3).

We used Integrative Genomics Viewer (IGV) (Thorvaldsdóttir et al., 2013) to visualize read alignments and remove false positive DNMs, defined as those missing in the genotypes due to local realignments and those in highly variable genomic regions (see Materials and Methods). On average, 17% of initial DNMs were removed as false positives (Tables S4-S6). To detect the false negative rate by simulation, we inserted 1,000 artificial DNMs to our read data. Our pipeline yielded an 89.5% detection rate, suggesting a 10.5% potential false negative rate, representing a good balance between Type I and Type II error.

### Distribution and variation of DNMs among families

After stringent variant filtering processes (see Materials and Methods), we observed a large variation in DNM rate across guppy pedigrees, with an average DNM rate of 0.60, 2.90 and 0.56 × 10^−8^ per nucleotide per generation in Fam1, Fam2 and Fam3, respectively (Table 1 & S1, Fig. 1a). The average mutation rate in Fam2 is significantly different from those in Fam1 (Wilcoxon rank-sum test, *P* = 2 × 10^−4^) and Fam3 (Wilcoxon rank-sum test, *P* = 2 × 10^−4^) (Table 1 & Fig. 1a).

**Fig. 1.**
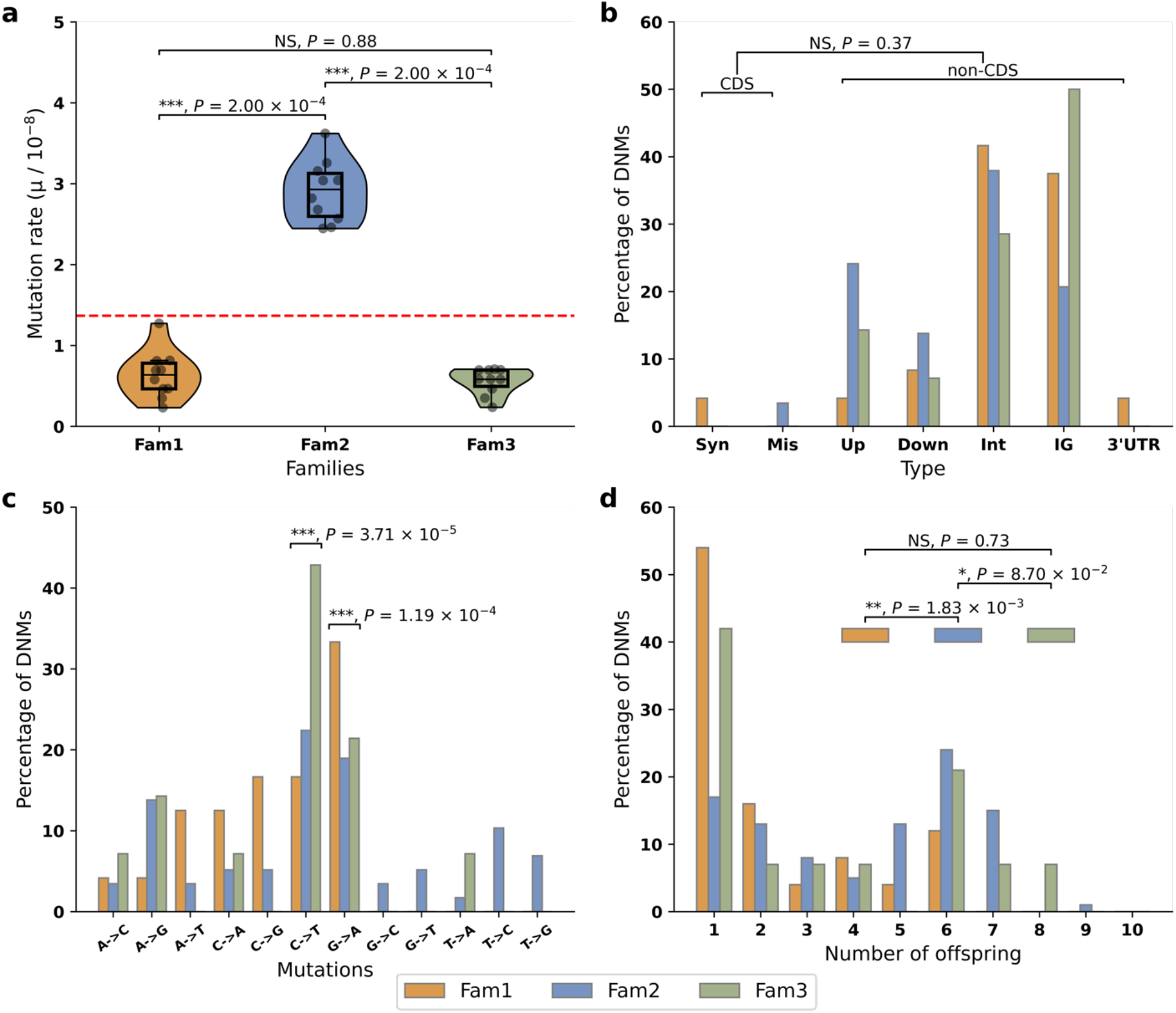
a. Mutation rate in three families. Red line represents the average mutation rate across three families. Stars indicate statistical difference among families. (Wilcoxon signed rank test, *** P < 0.001; NS: non-significant). Orange: Fam1; blue: Fam2; green: Fam3. **b. Functional distribution of DNMs**. There was no statistical difference (Fisher’s exact test, NS: non-significant) in the proportion of DNMs between coding sequences (CDS), including synonymous mutations (Syn) and missense mutations (Mis), and non-coding sequence (non-CDS), including upstream (Up), downstream (Down), intron (Int), intergenic (IG), and 3’ UTR variant (3’ UTR) once correcting for callable genome size. **c. Mutation spectrum**. Stars on the top of each category indicate significant difference to other categories (χ^2^ test, *** *P* < 0.001). **d. Shared mutations among siblings**. 45.73%, 72.76% and 57.14% of DNMs are shared by at least two siblings in Fam1, Fam2 and Fam3 respectively, likely reflecting mutations occurring during early germline development. DNMs observed in only a single offspring represent 54.17%, 17.24% and 42.86% of all DNMs in Fam1, Fam 2 and Fam3 respectively, and these most likely represent mutations after the onset of gametogenesis (Goldmann et al., 2019). The distribution of DNMs in a single offspring and DNMs shared by multiple siblings in Fam2 is significantly different from other two families (χ^2^ test, ** *P* < 0.01; * *P* < 0.1; NS: non-significant).

Even though the proportion of DNMs in coding sequence was far lower than those in non-coding sequence (Fig. 1b), this was non-significant after correcting for vast differences in callable genome size between these categories (Fisher’s exact test, *P* = 0.38, Fig. 1b). We observed significantly more transitions associate with CpG sites, including C > T transitions (χ^2^ test, adjusted *P* = 3.71 × 10^−5^, Fig. 1c, Table S7) and the reverse complement G > A transitions (χ^2^ test, adjusted *P* = 1.19 × 10^−4^, Fig. 1c, Table S7), consistent with findings in other species (Goldmann et al., 2016; Jónsson et al., 2017; Kessler et al., 2020). However, there is no significant difference in mutation spectra across families (χ^2^ test, *P* = 0.20, Fig. 1c).

Theoretically, DNMs can occur at any stage through development as well as through the process of gametogenesis itself. Developmental DNMs would be expected to affect a larger proportion of gametes, and would be expected to be shared among multiple siblings. DNMs occurring later, or in gametogenesis itself, would result in a small proportion of affected gametes and would not be expected to be shared across multiple siblings. Importantly, averaging across all our families, less than half of DNMs are observed in single offspring (Fig. 1d, Fig 2, Tables S1, S4-S6, S8).

**Fig. 2.**
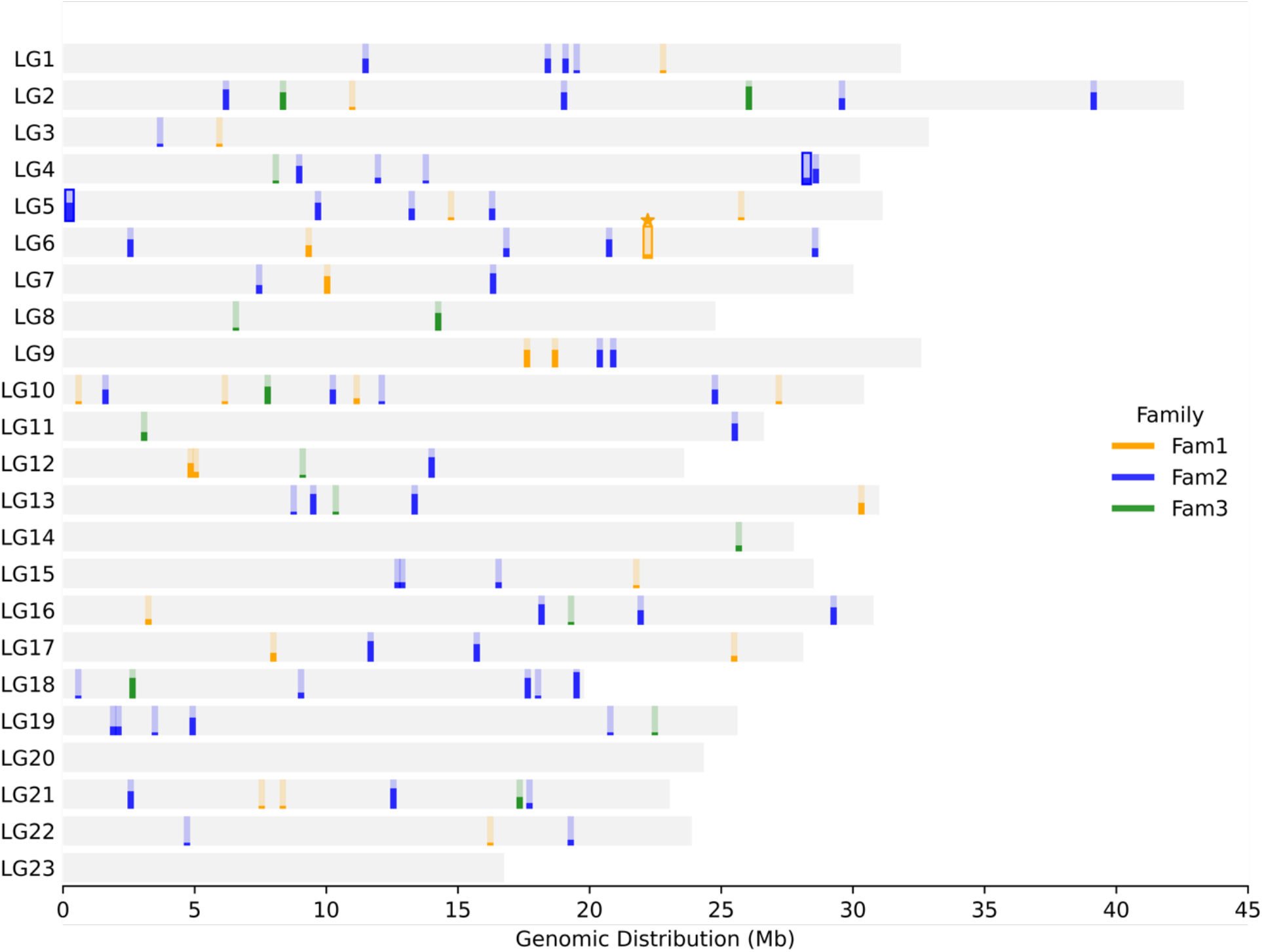
Genomic distribution of DNMs in three guppy families for each chromosomal linkage group (LG). The height of each bar represents the number of offspring sharing the same DNM in the family. DNMs with an outline represent those in coding regions. Stars (★) indicate synonymous mutations. No chromosome exhibits significant deviation from the expected number of DNMs based on its proportion of callable sites (χ^2^ test, see details in Table S6). Fam1 (orange), Fam2 (blue) and Fam3 (green).

This high proportion of DNMs shared by at least two siblings suggests that many DNMs occur before the onset of gametogenesis, and result from replication errors during germline development (Goldmann et al., 2019). Importantly, Fam2 has a higher proportion of shared DNMs than either Fam1 (χ^2^ test, *P* = 1.83 × 10^−3^) or Fam3 (χ^2^ test, *P* = 8.70 × 10^−2^), suggesting that the higher overall DNM rate in this family is largely the product of more early developmental mutations.

The greater number of cell divisions associated with spermatogenesis is expected in many species to result in a male-mutation bias (Ellegren 2007), although we might expect male-biased mutation to be relatively low in guppies due to their short overall generation time (Reznick et al. 1997). To estimate male mutation bias in our guppy families, we phased each individual and matched offspring haplotype blocks bearing DNMs to each parent haplotype to determine parent of origin, omitting DNMs with ambiguous phasing. Our data shows a consistent male-to-female mutation rate ratio across all three guppy families of ≈1.5 (Table S4-S6, S9). Germline mutation rates can vary across the genome, depending on contextual genetic background (Carlson et al., 2018). The DNMs we observe are statistically random in terms of genomic distribution (Fig. 2, χ2 test, see Table S3)

## Discussion

Germline DNMs are passed on from parent to offspring, and represent a major source of genetic variation upon which evolution acts. As such, the rate of germline DNMs is critical for defining the adaptive potential and mutational burden of a population. Here we assessed three large pedigrees for DNMs in the guppy, a longstanding model of rapid ecological adaptation. Guppies can rapidly adapt to shifting ecologies (Reznick et al., 1990), and this raises important questions about whether selection acts in these cases on DNMs or on standing genetic variation. The role of DNMs in guppy adaptation has important implications to the molecular signatures we might expect to detect, namely hard sweeps associated with recent DNMs versus soft sweeps acting on standing genetic variation (Harris et al., 2018; Hermisson & Pennings, 2005, 2017).

Overall, our estimate of the germline DNM rate, 1.35 ×10^−8^, is roughly comparable to estimates of other vertebrate species (Fig 3; Table S10), suggesting that rapid DNMs may not be the primary locus of selection in rapidly adaptive guppy populations. Indeed, relatively few sites in the guppy genome exhibit evidence of the hard sweeps associated with recent DNMs (Fraser et al., 2015), and patterns are more consistent with soft sweeps on standing genetic variation (van der Zee et al., 2022).

**Fig. 3.**
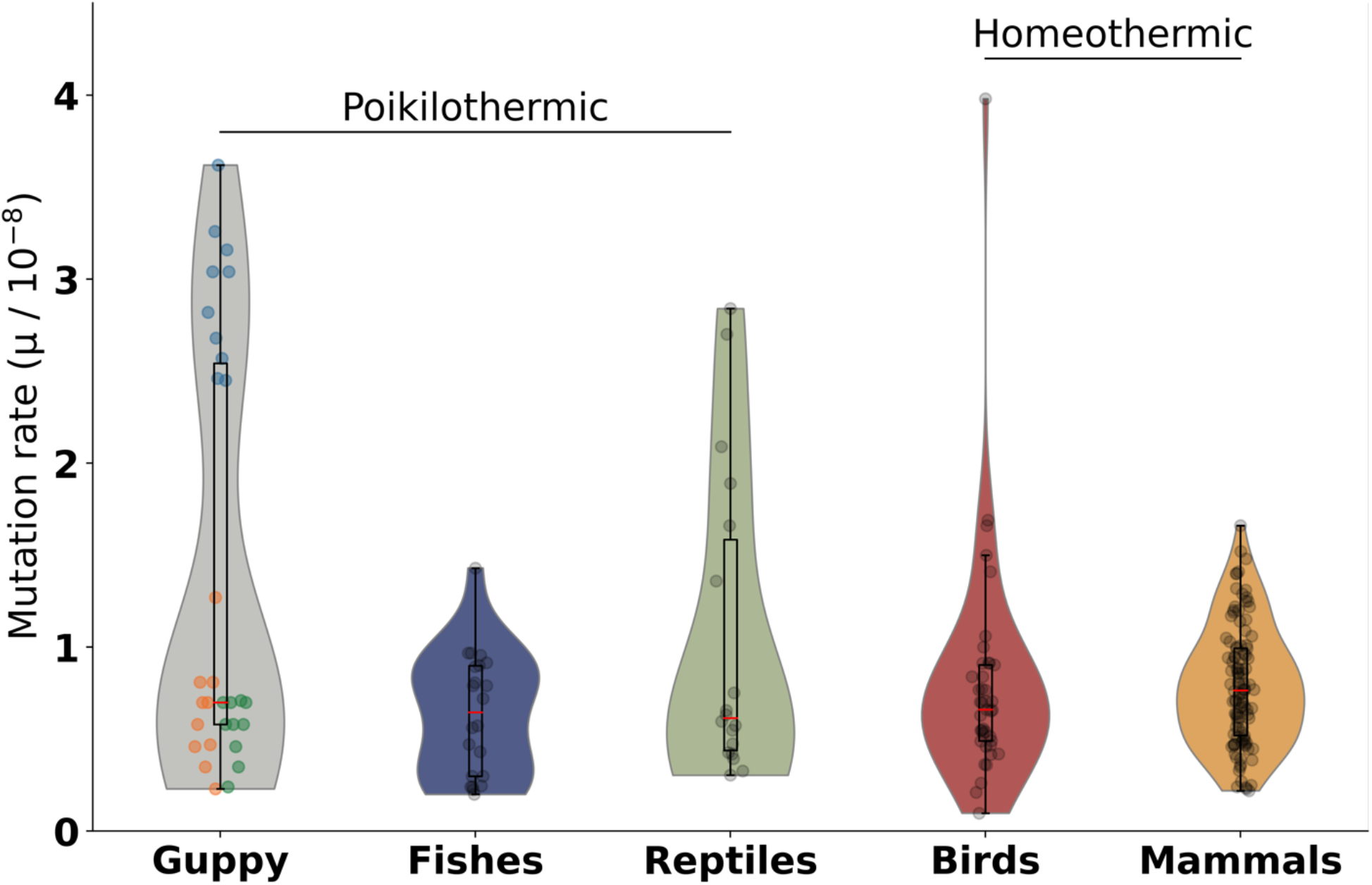
Distribution of trio DNMs in different animals. Distribution of trio DNM rate across different groups of poikilothermic animals, including fishes and reptiles, and homeothermic animals, including birds and mammals. Each data point represents a DNM rate in a parent-offspring trio. Three families in the guppy categories Fam1 (orange), Fam2 (blue) and Fam3 (green). More details could be found in Table S10. Datapoints from Awadalla et al. (2010); Bergeron et al. (2022, 2023); Besenbacher et al. (2015); Campbell et al. (2021); Conrad et al. (2011); Feng et al. (2017); Harland et al. (2017); Jónsson et al. (2017); Koch et al. (2019); Kong et al. (2012); Lindsay et al. (2019); H. C. Martin et al. (2018); Pfeifer (2017); Rahbari et al. (2016); Roach et al. (2010); Smeds et al. (2016); Tatsumoto et al. (2017); Thomas et al. (2018); Venn et al. (2014); Wang et al. (2020, 2022); Wu et al. (2020).

Our study design allows us to examine variation in the number of germline mutations across guppy individuals, and we observe a remarkable range in DNM rates across our families (Table 1). There is evidence of variation in germline mutation in some mammals, either as a result of increased paternal age (Carlson et al., 2018; Kong et al., 2012; Wang et al., 2022) or stochastic processes (Goldmann et al., 2021), however the degree of variation observed in these species is far less than we observe in guppies. Our data comes from captive reared guppies kept under controlled, consistent conditions, suggestion that the differences in DNM rate that observe are intrinsic, rather than due to environmental variation. Moreover, the short generation time (Reznick et al. 1997) and lifespan (Reznick et al., 2006) in guppies guarantees that the parents in our families varied very little in overall age, suggesting that paternal age may have little effect in this variation we observe.

More importantly in the context of guppies, the wide variation in DNM rate suggests that some lineages, with higher mutation rates, might in fact show more potential for adaption by DNMs than standing variation. Although overall, our results suggest that the rapid adaptation often observed in guppy populations (Gordon et al., 2015; Reznick et al., 1990; Reznick et al., 1997; Whiting et al., 2022) is largely the product of selection on standing genetic variation, the extensive variation in DNM rate we observe suggests that if heritable, DNM ratio might vary extensively in natural populations with concurrent variation in adaptive potential. The role of heritable variation in DNM rate and adaptive potential remains an important area for further exploration.

It is worth noting that alternative bioinformatic pipelines have yielded DNM rate estimates that vary by a factor of two for the same parent–offspring trios (Bergeron et al., 2022). This variation represents differences in the balance between stringent filtering to remove false positives, and the minimization of false negatives. Although we were stringent in our filtering and inclusion criteria, our simulations reveal that the pipeline yielded an 89.50% detection rate, suggesting a 10.5% potential false negative rate. Because we used the same pipeline across all families, any error from false negatives or false positives should affect all families similarly, allowing for direct comparisons.

Moreover, compared to other poikilothermic species, the absolute value of the mutation rate in the guppy is similar to whole genome pedigree-based estimates in poikilotherms, and also similar to rates seen in some endothermic primates (Fig 3, Table S10). The high DNM ratio in Fam2 largely explains the extensive range of DNM rate observed here (Fig. 1a, Table S4-S6), and without this family, our estimates would be similar to other fish species (Fig. 3). It might be expected that the lower metabolic rates in poikilothermic species would produce less oxidative damage to the DNA molecule (Adelman et al., 1988; A. P. Martin & Palumbit, 1993). However, poikilothermy does not itself explain DNM rate variation across vertebrates (Bergeron et al., 2023), suggesting that the role of homeostasis in DNM rate is small.

Our large pedigrees, each comprised of 10 offspring, make it possible to assess DNM sharing among siblings, and thus assess the degree of mosaicism in the germline genome (Goldmann et al., 2019). Our results suggest that the developmental timing of DNMs in the germline is highly variable (Fig. 1d). If a DNM occurs early in germline development, it will be passed on to a greater proportion of daughter cells than those later in development, and thus more likely to transmit to the next generations and be shared by multiple siblings. Although the number of shared mutations varied among our three families, ranging from 50% to 73%, it was nonetheless the majority of DNMs and indicates most inherited DNMs occur during germline development instead of after the onset of gametogenesis. This is consistent with studies in smaller human families which nonetheless revealed substantial sharing of DNMs among siblings and variation in the proportion among families (Rahbari et al., 2016).

We also assessed the male mutation bias in DNM generation. Male mutation bias occurs from the increased number of cell divisions in spermatogenesis compared to oogenesis (Ellegren, 2007). This male-biased mutation rate scales with increasing parental age (Carlson et al., 2018; Francioli et al., 2015; Kong et al., 2012), and the relatively short lifespan of guppies might suggest a minimal male mutation bias. Indeed, our estimate of ≈1.5 is on par with that of other short-lived species (Wilson Sayres & Makova, 2011).

Overall, our results indicate that the developmental timing, rate and spectrum of DNMs varies substantially across families in guppies, suggesting that mutation rate as a trait can vary across at the level of individuals, families, and populations and might be able to evolve under different conditions (Lynch et al., 2016). Our results also suggest that comparisons across species based on just one or a few parent-offspring trios may experience high error due to intra-specific variation. Taken together, our results present an important advance in our understanding of the variation in germline DNMs, and their contribution to adaption.

## Methods and Material

### Samples and DNA sequencing

We selected three families from a previous lab experiment (Morris et al., 2020), each of which contains one father, one mother, five sons and five daughters. In total, 36 samples from three pedigrees were collected. DNA was extracted from head tissues with Qiagen DNeasy Blood & Tissue Kit, following the manufacturer protocol. After DNA quality control, all the samples were sequenced individually on the Illumina NovaSeq 4000 platform.

### Reference genome reconstruction

Divergence between the individual of the Ensembl reference genome v.99 (Künstner et al., 2016) and our lab population might trigger false positive DNM calls and complicate downstream DNM detection. Our lab population originated from a high predation population of the Quare River in Trinidad. We therefore selected the most contiguous 10X Genomics Linked Read female genome assembly from the Quare high predation population, presented in Almeida et al. (2022).

We customized our reference genome first by assembling the genome using LongRanger v2.1.1 (10X Genomics), removing redundant short sequences for which 95% of contigs overlapped with longer ones. We further improved the assembly using a *K-mer* based approach, ARKS+LINKS v1.0.4 (Coombe et al., 2018) and anchored scaffolds into chromosomes using RAGTAG v2.1.0 (Alonge et al., 2019, 2022). Genome completeness was assessed by QUAST v5.1.0 (Mikheenko et al., 2018) and BUSCO v 5.3.2 (Simão et al., 2015), see details in Table S11 & S12.

We performed *de novo* prediction and modelling of transposable element (TE) families using RepeatModeler2 (Flynn et al., 2020) with default parameters in both the Ensembl reference genome and the reconstructed genome. Repetitive sequences were then identified by RepeatMasker v.4.1.1 (Smit, 1996) using databases outputted by RepeatModeler2 (Flynn et al., 2020). In total, 27.04% of sequences were identified as repetitive sequences and thus were removed (See details in Table S13).

### Alignment, Genotyping and SNP Filtering for DNMs detection

We used FastQC v0.11 (Andrews S, 2018) and Trimmomatic v0.36 (Bolger et al., 2014) to remove adapter sequences and low-quality reads. After quality control, we recovered ∼25x average read depth per sample. High-quality reads were aligned to the reconstructed female genome, using BWA 0.7.15 MEM (Li & Durbin, 2009) with default parameters. We filled in mate coordinates and mate related flags, sorted alignment by coordinates, and marked PCR duplicates with Picard.

To reduce the chance of removing true DNMs in each individual, we conducted parent-offspring trio-based genotyping instead of the joint genotyping with all offspring in a family. We called genotypes with two independent haplotype-aware software, GATK4 and BCFtools, respectively, using only those sites called by both approaches (see details in Table S2).

First, we applied hard filtering to the raw SNP dataset in trios following the GATK best practice ’QD < 2.0, FS > 60.0, SOR > 3.0, MQ < 40.0, MQRankSum < -12.5 and ReadPosRankSum < - 8.0’ for raw GATK4 genotype dataset and ’MQBZ < -3 || RPBZ < -3 || RPBZ > 3 || FORMAT/SP > 32 || SCBZ > 3 || TYPE=“INDEL”’ for raw BCFtools genotype dataset. We selected biallelic SNPs in which genotypes are homozygous in parents but heterozygous in offspring with Mendelian violation quality score > 30 using GATK ’SelectVariants’, and restricted SNPs with genotype quality > 30 and with no genotype-level missing data included for further analysis. The difficulty in definitively mapping reads from repetitive regions makes it impossible to assess DNMs in these regions, therefore we also excluded any SNPs that fall in the genomic regions identified as low complexity genomic regions by RepeatModeler2 (Flynn et al., 2020) and RepeatMasker v.4.1.1 (Smit, 1996) (see *3*.*7 Repetitive Genomic Regions Identification*).

To further limit the false DNMs discovery rate, we filtered SNPs at the individual level as well. We removed all sites where coverage between < ½ X or > 2X of individual average sequencing depth. For each trio, all three family members were required to pass these criteria for a site to be retained. Variant callers such as GATK4 and BCFTools determine individual genotypes based on calculated genotype likelihood. This in fact affects DNM calls as the false discovery rate can arise when heterozygous loci of parents are miscalled as homozygous. Thus, each parental genotype was required to have alternative allele depth to be zero. For heterozygous loci in the offspring, we performed a 2-sided binomial test, under the null hypothesis of 0.5 relative frequency of reads supporting reference allele and alternative alleles, with a cut-off *P-value* of 0.05, following methods in Wu et al., (2020) (Table S2).

SNPs that are in genomic regions with complex read alignments would be problematic for DNMs detection and usually indicate misalignments. Following methods in Wang et al., (2022), we further filtered any DNMs located in genomic regions where >50 % of aligned reads contain gaps or are multiply mapped to other genomic locations.

Finally, we visualized each inferred DNM in IGV, an interactive tool for the visual exploration of genomic data. We removed inferred DNMs where the DNM was observed in the parent, but were removed during the realignment step in GATK4 and BCFtools, which can exclude alternative alleles.

### Kinship Coefficient Analysis

To verify parentage in our pedigrees, we first called genotypes across each family using the GATK4 ’HaplotypeCaller’ with the following parameters: ’--linked-de-bruijn-graph true --min-pruning 0 --recover-all-dangling-branches true’ in ’-ERC GVCF’ mode. Then, we genotyped across individuals in each pedigree using GATK4 ’CombineGVCFs’ and ’GenotypeGVCFs’ with default parameters. The pairwise kinship coefficient in each family was calculated using KING (Manichaikul et al., 2010) with default parameters.

### Mutation Rate and Callable Genome Size Estimation

To calculate the mutation rate, we estimated the callable genome size using GATK4 ’HaplotypeCaller’ with the following parameters: ’--linked-de-bruijn-graph true --min-pruning 0 --recover-all-dangling-branches true’ in ’-ERC BP_SOLUTION’ mode in each individual. For each individual, we removed sites with coverage that deviated from average individual sequencing read depth, and sites located in the repetitive genomic regions predicted as described above for filtering DNM sites. Lastly, SNPs that were heterozygous in either parent were excluded from the callable genome size estimation. The final callable genome size was intersected across each trio (Table S3).

Mutation rate calculation followed the equation below, where μ is the mutation rate, M_*i*_ is *i*^*th*^DNM and C_*i*_ is the *i*^*th*^ intersected callable genome loci of each trio.

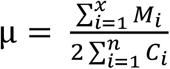

### Genotype Phasing and Male Mutation Bias Estimation

To understand paternal and maternal effects on DNMs, we first phased each individual using WhatsHap in read-based phasing mode with default parameters (Martin et al., 2016). Phasing blocks in offspring bearing DNMs were then matched back to the parent’s phasing block to determine DNM phasing results. DNMs with ambiguous phasing were left unphased. Finally, we visualized read alignment from trios using IGV to check the linkage of DNMs and adjacent variants and validate genotype phasing results and inferred the parent of origin for phased DNMs.

### Mutational Spectra Identification

Based on SnpEff (Cingolani et al., 2012) and the guppy reference genome v.99 (Künstner et al., 2016), we were able to annotate each variant and characterize the mutational spectra of DNMs. To test whether the number of DNMs located in coding regions are significantly different than expected based on the coding callable genome, we counted the total number of loci in coding regions and non-coding regions in callable genome size, and employed Fisher’s exact test.

### Simulation of DNM Detection Pipeline

To determine the false negative rate of our DNMs detection pipeline, we employed a simulation approach by inserting artificial mutations to the aligned reads of the offspring using BAMSurgeon (Ewing et al., 2015). We randomly simulated 1000 DNMs and restricted the insertion sites to the region of callable genomic region as described above (*Mutation Rate and Callable Genome Size Estimation*) and non-variable sites where less than half of reads contain indels and gaps, as described in previous studies (Campbell et al., 2021; Wu et al., 2020). Then, we went thought the full pipeline (Fig. S4) to detect the number of DNMs and determined the proportion of simulated DNMs that we detected.

## Data availability

Raw sequence data will be deposited to NCBI BioProject XXXXX.

## Code availability

The code used for processing the data will be deposited at XXXX (GitHub) upon manuscript acceptance.

## Acknowledgements

This work was funded by NSERC and a Canada 150 Research Chair (to JEM) and a Doctoral Scholarship from the China Scholarship Council (to YL). We thank Tom Booker for helpful suggestions on the simulation and Pedro Almeida for the Quare reference genome assembly. We also thank members of the Mank Lab for helpful feedback through the course of this project, as well as constructive comments on a previous version of this manuscript.

## Author contributions

JEM and YL conceived the study and designed the experiments and analysis. ID, JM and JEM collected the data. YL and ID performed DNA extractions. YL, WvdB, and ID performed the data analysis. JEM and YL wrote the manuscript and all authors contributed to revisions.

## Supplementary material

**Supplementary Figure 1.**
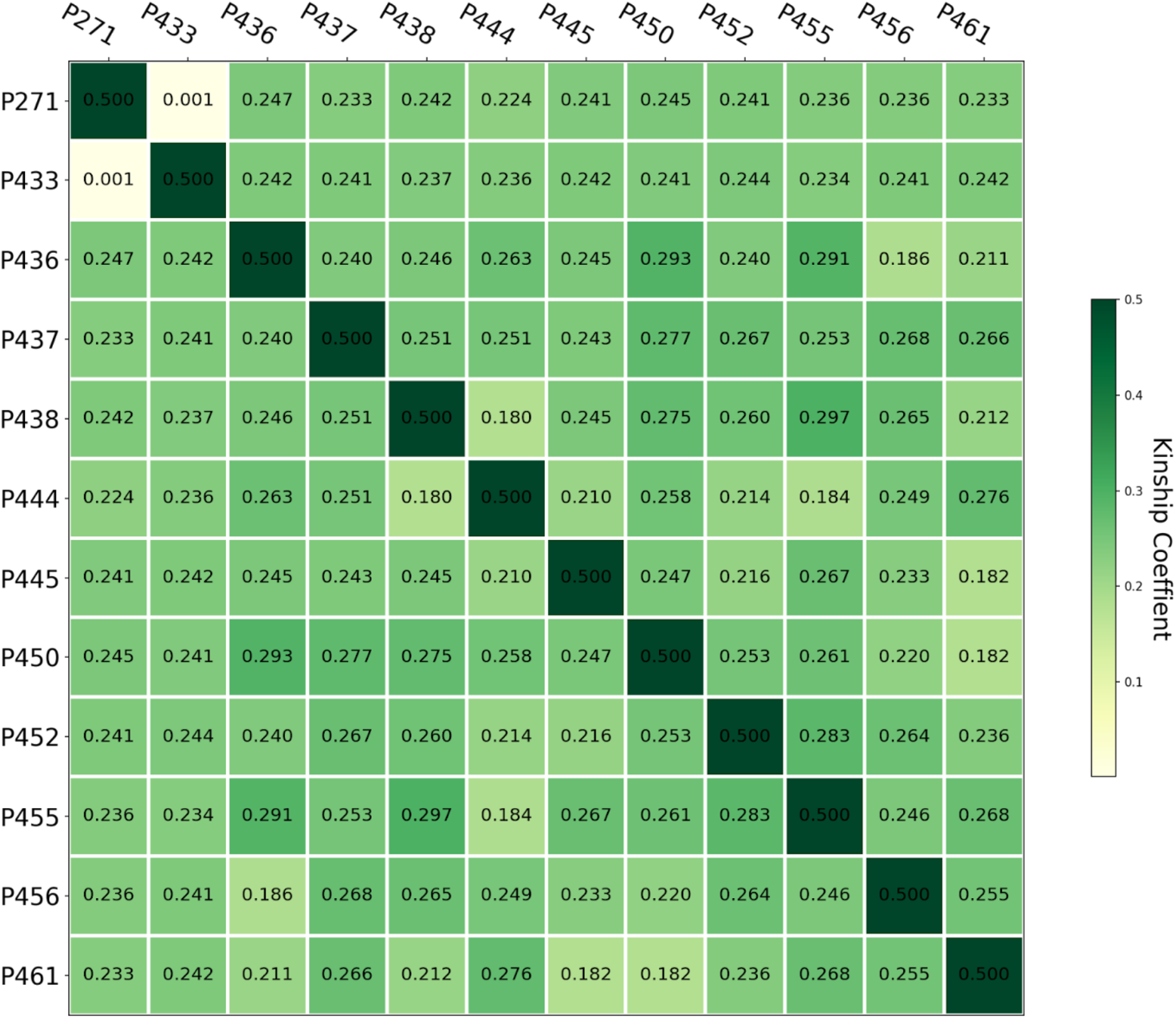
Kinship coefficient of Family 1. Father: P271; Mother: P433; other individuals are offspring in this family. Kinship coefficients among siblings indicate second degree relationship and kinship coefficients between parents indicates an unrelated relationship.

**Supplementary Figure 2.**
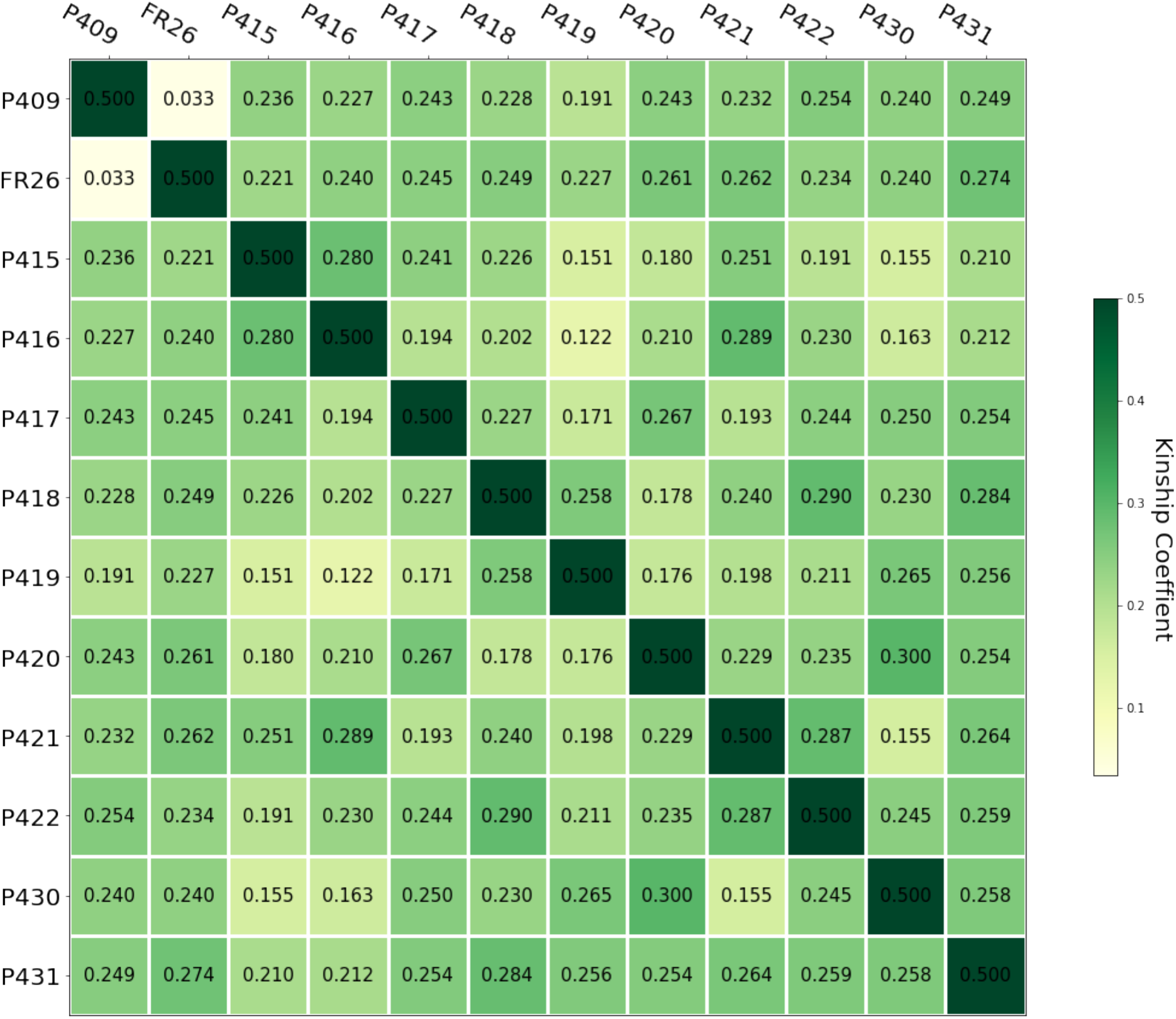
Kinship coefficient of Family 2. Father: P409, Mother: FR26; other individuals are offspring in this family. Kinship coefficients among siblings indicate second degree relationship and kinship coefficients between parents indicates an unrelated relationship.

**Supplementary Figure 3.**
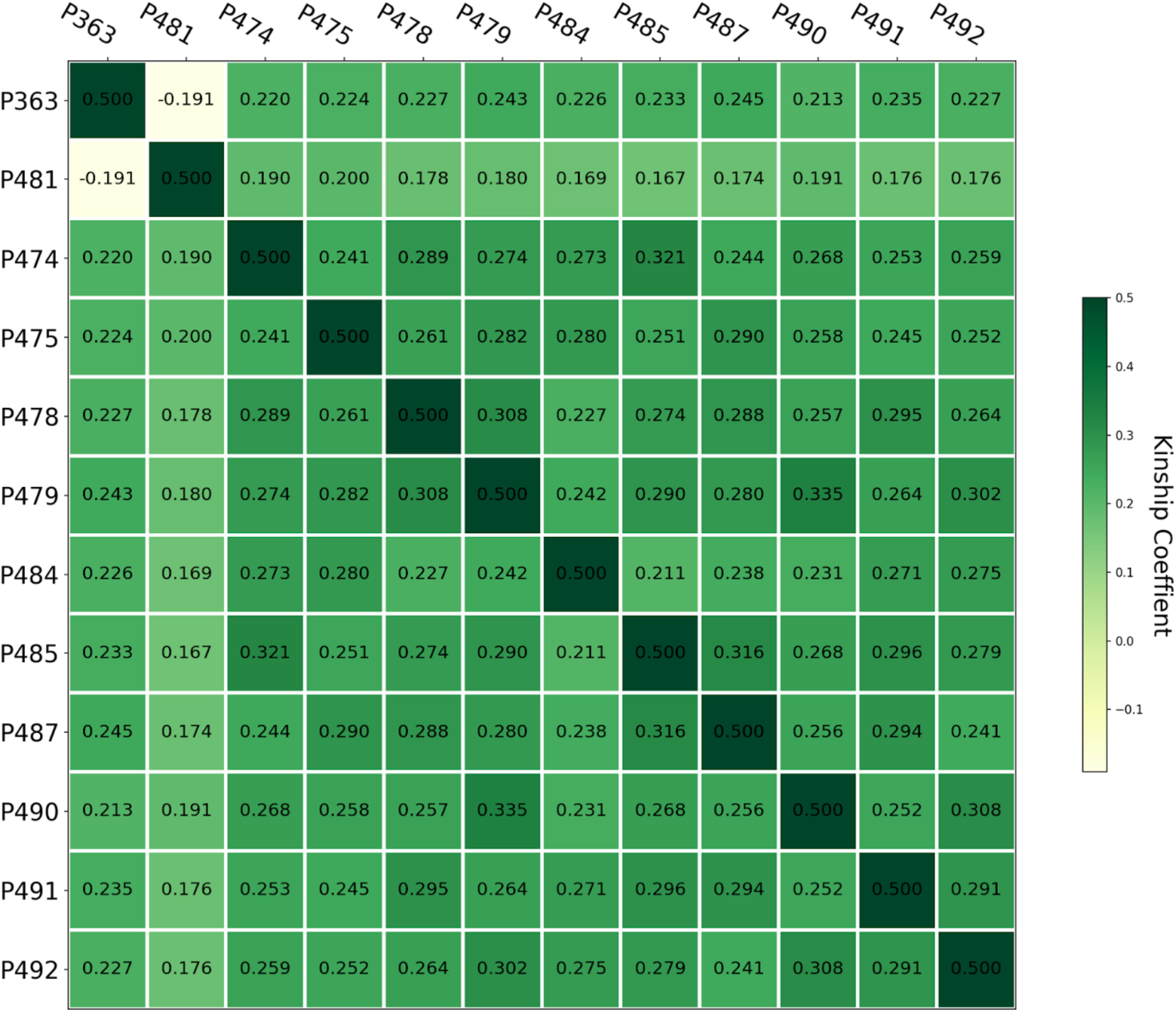
Kinship coefficient of Family 3. Father: P363, Mother: P481; other individuals are offspring in this family. Kinship coefficients among siblings indicate second degree relationship and kinship coefficients between parents indicates an unrelated relationship.

**Supplementary Figure 4.**
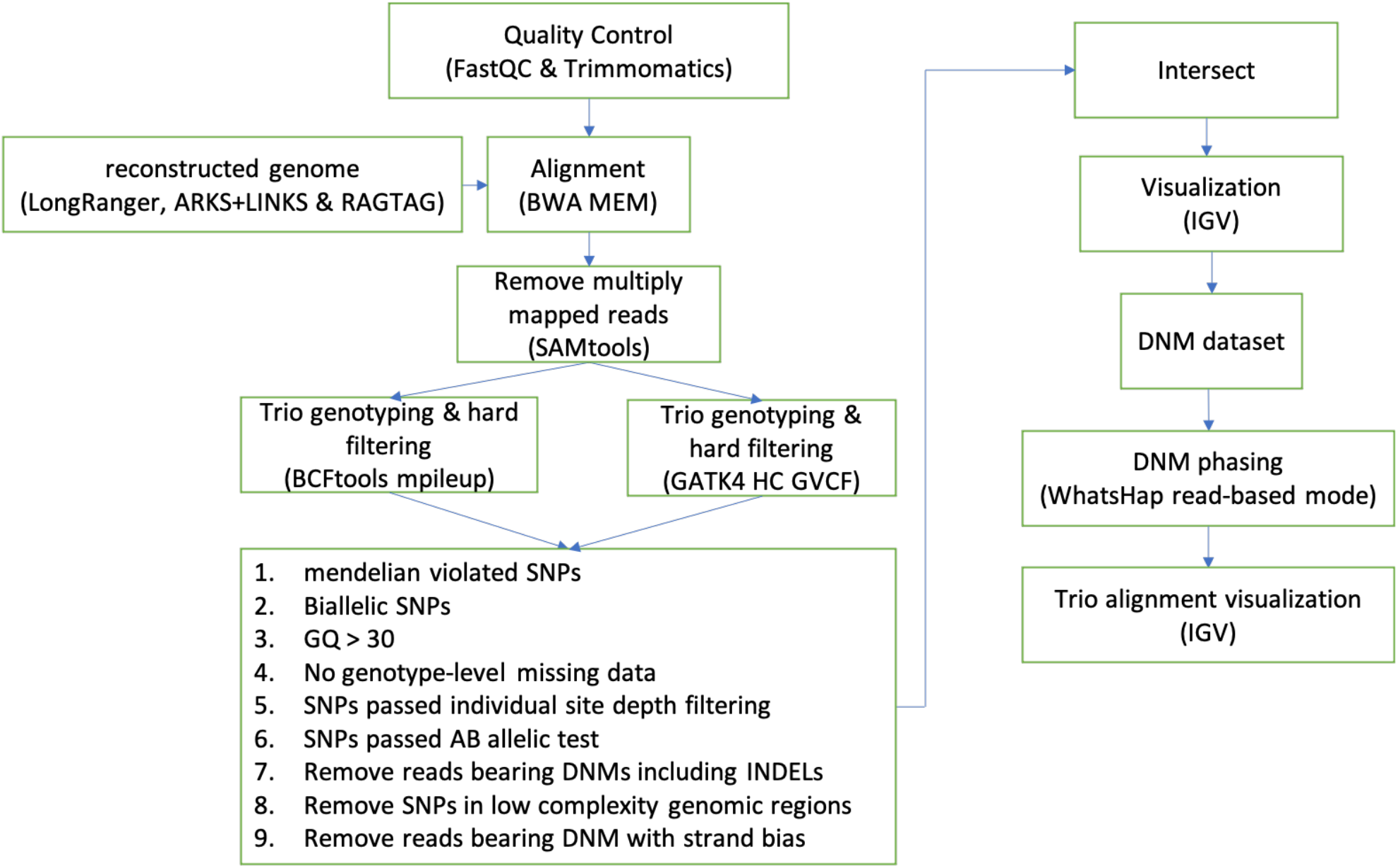
DNM detection pipeline. The software for each step is indicated in the brackets. Details are in Material and Methods

**Supplemental Table 1. Summary statistics of each sample in three guppy pedigrees**

**Supplemental Table 2. Summary Statistics of variant filtering in the pipeline**

**Supplemental Table 3. Callable genome size of each individual**

**Supplemental Table 4. Summary Statistics of DNMs in Fam1**

**Supplemental Table 5. Summary Statistics of DNMs in Fam2**

**Supplemental Table 6. Summary Statistics of DNMs in Fam3**

**Supplemental Table 7. Mutation spectra in three families**

**Supplemental Table 8. Number of offspring sharing DNMs in each family**

**Supplemental Table 9. Parental mutation ratio**

**Supplemental Table 10. Germline mutation rate of poikilothermic and homeothermic vertebrate species using whole genome pedigree sequencing**

**Supplemental Table 11. Summary statistics of reconstructed genome with BUSCO**

**Supplemental Table 12. Summary statistics of reconstructed genome with QUAST**

**Supplemental Table 13. Result of RepeatMasker with reconstructed genome**

